# Fully Transient CRISPR/Cas12f system in plants capable of broad-spectrum resistance against Begomovirus

**DOI:** 10.1101/2022.06.07.495110

**Authors:** Sibtain Haider, Ali Faiq, Muhammad Zuhaib Khan, Shahid Mansoor, Imran Amin

## Abstract

CRISPR/Cas system has emerged as the most efficient genome editing technology for both eukaryotic and prokaryotic cells. Recently, biologist have started using CRISPR/Cas9 as a defence machinery in plants against DNA viruses by targeting conserve regions of their genome. Considerable resistance requires formation of stable transgenic lines with multiple gRNAs, targeting specific viruses. Development of such transgenic plants is not only time consuming but also there will always be some uncertainty of their efficiency and off targets in plant genome. Newly discovered miniature CRISPR/Cas12f (Cas14a) system has unique ability to chop down every ssDNA fragment once activated through targeted cleavage. Here we report first fully functional transient CRISPR/Cas12f system in plants. We also show that Cas12f with just one gRNA is enough for substantial broad-spectrum resistance against Gemini viruses. Plant phenotype showed nearly complete recovery and qPCR results showed multifold decrease infection of CLCuV in CRISPR/Cas12f treated plants compare to the infected plants (infected with CLCuMuV and CLCuKV). Smaller size, broad range and more efficiency make Cas12f a superior alternative of Cas9 against diverse group of ssDNA viruses.

## Introduction

Begomoviruses are circular ssDNA plant viruses. They belong to the family of *Geminiviridae* with a genome size of nearly 2.8 kb (monopartite) to 5.4 kb (bipartite). The principal vector is whitefly (*Bemisia tabaci*) and responsible for serious damage to many economically important crops. One of the most devastating group is Cotton leaf curl viruses (CLCuV) and the major constraint to cotton production across Pakistan and India. These viruses cause cotton leaf curl disease (CLCuD) characterized by vein thickening and leaf curling. All of CLCuVs are monopartite except cotton leaf crumple virus [1]. In the early 1990s during the first epidemic of cotton, production was devastated in Pakistan due to Cotton leaf curl virus now known as “Multan strain”. Soon resistant varieties of cotton were introduced in the field but in 2001, these resistant varieties started to show symptoms similar to CLCuD. Later, this leads to the second pandemic in the cotton crop by a mutant strain of CLCuV now known as “Burewala strain”. It has been hypothesized that CLCuD can spread to other cotton-producing areas of the world where there is no sign of this disease now and recent occurrence of CLCuD in china has supported this hypothesis.

The genome of monopartite begomovirus contains six open reading frames (ORF) with genes V1 and V2 in the sense direction. V1 encodes for coat protein (CP), V2 is supposed to be responsible for cell-to-cell movement and repressing cellular RNAi mechanism. While in complementary sense direction, Virus encodes four proteins C1, C2, C3, C4. C1 which are responsible for replication protein (Rep), C2 codes for transcription activation protein (TrAP), C3 codes for replication enhancer protein (REn) and C4 is responsible for the repression of host’s RNAi mechanism. Begomoviruses also have non-coding region known as intergenic region (IR). This region contains a bidirectional promoter and highly conserve nona-nucleotide sequence (TAATATTAC) [2, 3].

Management of Begomovirus based diseases is challenging and expensive due to their diversity and high mutation rate. Different approaches have been used to minimize the infection including use of pesticide against whitefly vector or use of resistant varieties in the field. RNAi based approaches have also been used to target the golden mosaic virus (BGMV) [4] [5–7]. However, these approaches are good for some closely related viruses while in field condition multiple begomovirus species effects on plant and effectiveness of these approaches decline. An efficient, robust and highly durable strategy is required to tackle this menace.

CRISPR/Cas system is a bacterial defence mechanism against viral particles and invading plasmid but recently it has arisen as a most efficient programable genome editing tool. Unlike its predecessors like Zinc Finger Nuclease (ZFN) and Transcription Activator-like Effector Nucleases (TALENs), CRISPR/Cas is much more diverse, programable, robust and target-specific. Based on great diversity, the CRISPR/Cas system has been divided into 35 families. Initially researchers used CRISPR/Cas9 to induce targeted double standard DNA break in plant genome and incorporate desirable characteristics. Shortly, it was realized that the CRISPR/Cas system can also be used as a defence mechanism against plant viruses. The Cas9 nuclease was exploited for this purpose against plant DNA viruses (*Geminiviruses*), because of its ability to efficiently recognise and produce dsDNA breakage. This dsDNA breakage results in either DNA degradation or repaired by error-prone non-homologous end joining (NHEJ) in viruses. NHEJ mostly cause drastic mutations in the viral genome leading to non-translated products. *Ali* et al demonstrated the efficiency of CRISPR-Cas9 machinery against coding and non-coding sequences. They observed that targeting coding region may result in a mutated virus which will not be prone for Cas9 derive immunity [8–11].

Similar results were obtained when Cas9 was used in Cassava against African cassava mosaic virus (ACMV). Single gRNA was inadequate to confer resistance and it increased virus mutation rate [12]. A Roy, *et al* has reported that successful defence against Gemini-virus requires multiple gRNAs targeting conserve region of virus [13]. Even though results are improving with altered approaches but still there are some drawbacks associated with the use of Cas9. Most importantly the size of Cas9, efficient regulation of such big protein with multiple gRNAs in transgenic lines is very difficult. Another problem with Cas9 is a requirement of PAM sequence, which limits its target regions, and possibility of off targets in host genome. Immunity due to CRISPR/Cas9 display some disadvantages as with other approaches.

LB Harrington., *et al* has discovered a new CRISPR/Cas system consist of a very small Cas protein (Cas12f) and a relatively long gRNA [14]. Initially, it was suggested that it can only cut ssDNA and does not require any PAM sequence. T Karvelis., *et al* has proved that Cas 14a can also recognise T-rich PAM sequence to cut dsDNA and they edited genes in the human cell through Cas12f [15]. This makes Cas12f a very valuable tool against both ssDNA and dsDNA targets. The Cas12f can be a great tool to target ssDNA viruses in plants and will have superiority over Cas9 because of its ability to do collateral cleavage of ssDNA once activated. [16].

In plants, development of transgenic lines is a long tedious process there will always be some uncertainty of their efficiency and off-targets in the plant genome. Transient expression provides the scientist with a better alternative to test the efficacy of their system. Plant viral vectors have been used for the expression of commercial proteins and can efficiently transiently express foreign proteins. Diverse and efficient genomic machinery make viral vectors an excellent choice for transient systemic expression. Different groups have tried to make transient CRISPR/Cas system using viral vectors but the large size of Cas9/Cas12/Cas13 proteins make it problematic to create a complete transient system. However, the smaller size of Cas12f provides a possibility of transient expression through RNA viruses. Here, we have developed first fully transient CRISPR/Cas system capable of inducing immunity against ssDNA viruses in plants. A combination of RNA viruses Potato virus X (PVX) and Tobacco rattle virus (TRV) is used to express Cas12f and gRNA respectively [17–19]. Both components of CRISPR/Cas12f system were expressed successfully. Co-inoculation of these components was cable to induce high immunity against a broad range of plant ssDNA virus by using only a single gRNA.

## Results

Sequencing results from CLCuD infected samples revealed high prevalence of CLCuMuV was identified from Vehari, Pakistan with multiple begomoviruses. Phylogenetic tree and complete genome analysis of has been published by our lab [20]. Multiple sequence alignment was used to identify most conserve region (precoat protein).

The gRNA cassette against the precoat protein of CLCuMuV was cloned in binary vector TRV-gRNA. While the Cas12f gene was cloned in pgR106 and named as pgR-Cas12f (table 2). Both constructs were confirmed with restriction digestion analysis, sequencing and were agroinfiltrated in *N. Benthamian* for expression analysis. Plant showed mild symptoms of TRV and PVX respectively.

**Table 1.** Inoculation

|  | Multan Virus (MV)<br>Infectious clone | Kokran Virus (KV)<br>Infectious clone | pgR-Cas12f | TRV2-gRNA | pTRV-RNA1 |
| --- | --- | --- | --- | --- | --- |
|  |  |  |  | TRV-gRNA |  |
| Healthy Control | - | - | - | - | - |
| CLCuMuV<br>positive | + | - | - | - | - |
| CLCuKV positive | - | + | - | - | - |
| Cas12f Expression | - | - | + | - | - |
| gRNA Expression | - | - | - | + | + |
| Cas12f-CLCuMuV | + | - | + | + | + |
| Cas12f CLCuV<br>(K+M) | + | + | + | + | + |

\*All the plasmids were transformed in *Agrobacterium* strain GV101. Equal volumes of each inoculum were mixed to obtain a final volume of 20 ml and 2 ml inoculum per plant was used.

**Table 2.**
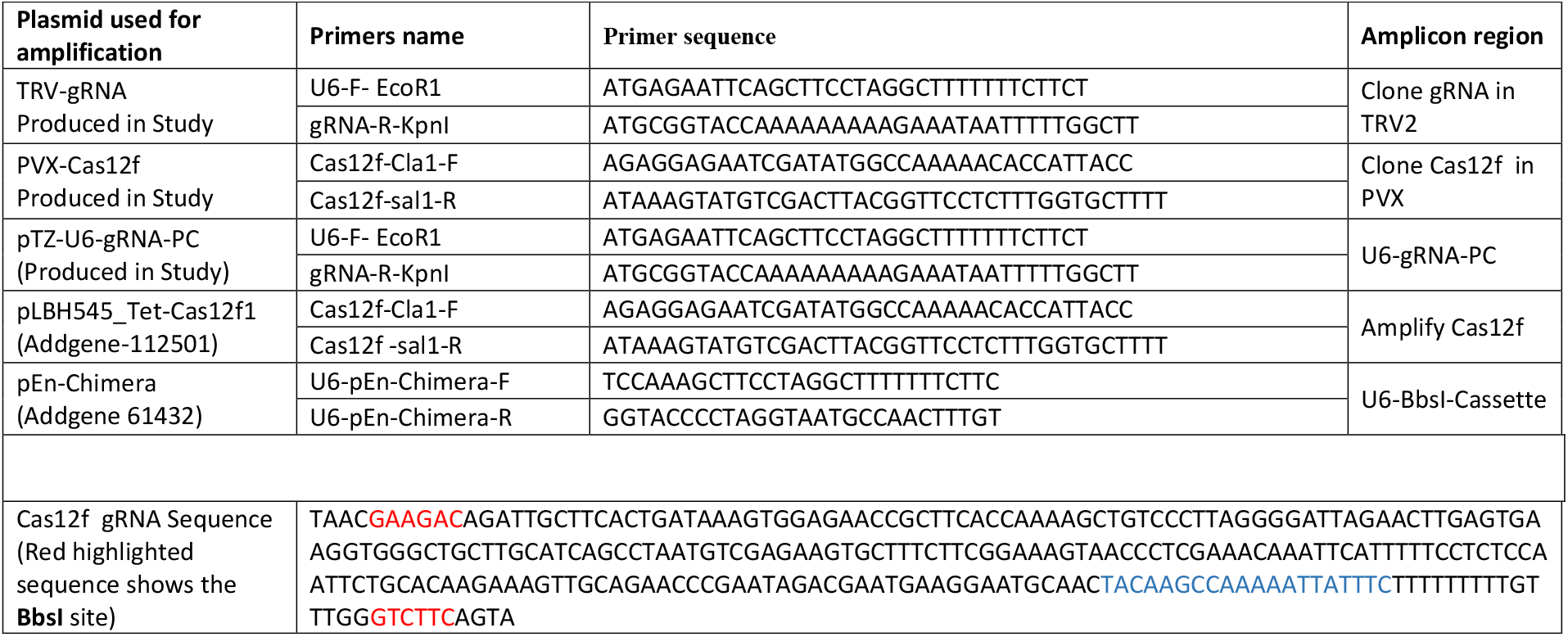
Primers and gRNA sequences

DNA was isolated from plant leaves at 15dpi, RT-PCR showed full-length systemic expression of Cas12f and gRNA cassette. Results were encouraging and we decided to inspect if Cas12f and gRNA can be used to target and digest the ssDNA or in our case plant ssDNA viruses. To determine the effectivity of Cas12f against CLCuMuV plants were agroinfiltrated with a mixture of pgR-Cas12f, TRV-gRNA and infectious clone of CLCuMuV. These plants showed very few symptoms of CLCuD after two weeks and almost zero symptoms after 4 weeks of inoculation. While control plants with only infectious clones of CLCuMuV showed severe symptoms of CLCuD even after 2 weeks of inoculation (Figure 3).

**Figure 1:**
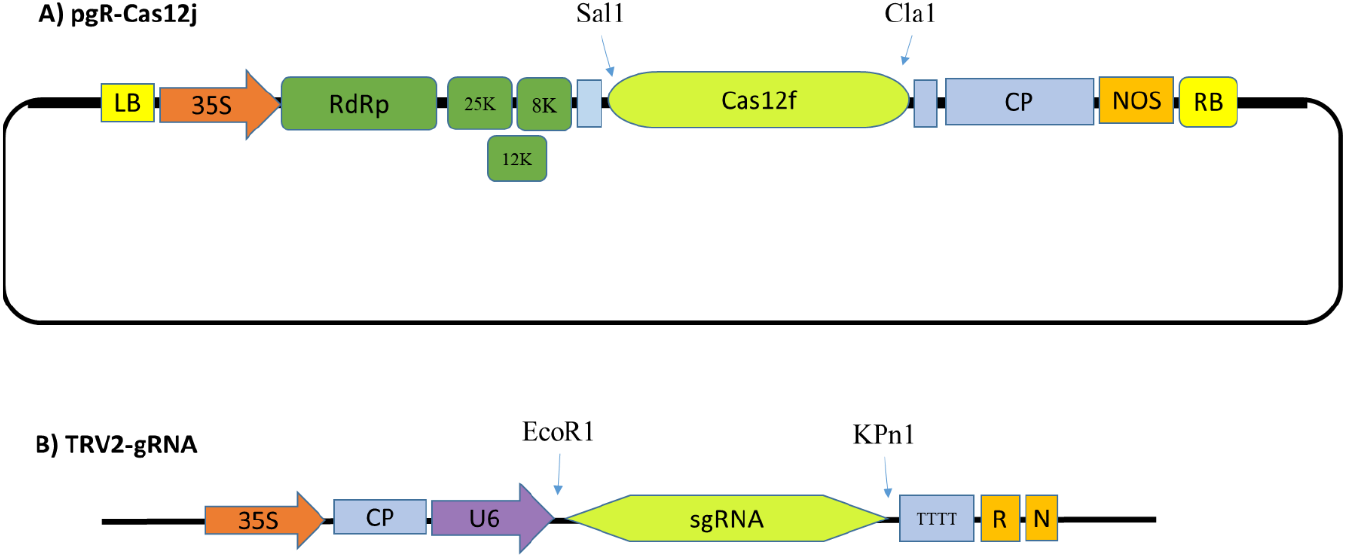
vector maps of pgR-Cas12f and TRV2-gRNA. **(A)** Confirmed PCR amplicon of Cas12f was cloned in pgR106 vector at Cla1 and Sal1 site to create pgR-Cas12f. **(B)** The U6-gRNA-pc cassette was cloned in pTRV-RNA2 at EcoR1, Kpn1 restriction site and the resultant vector was called TRV2-gRNA.

**Figure 2:**
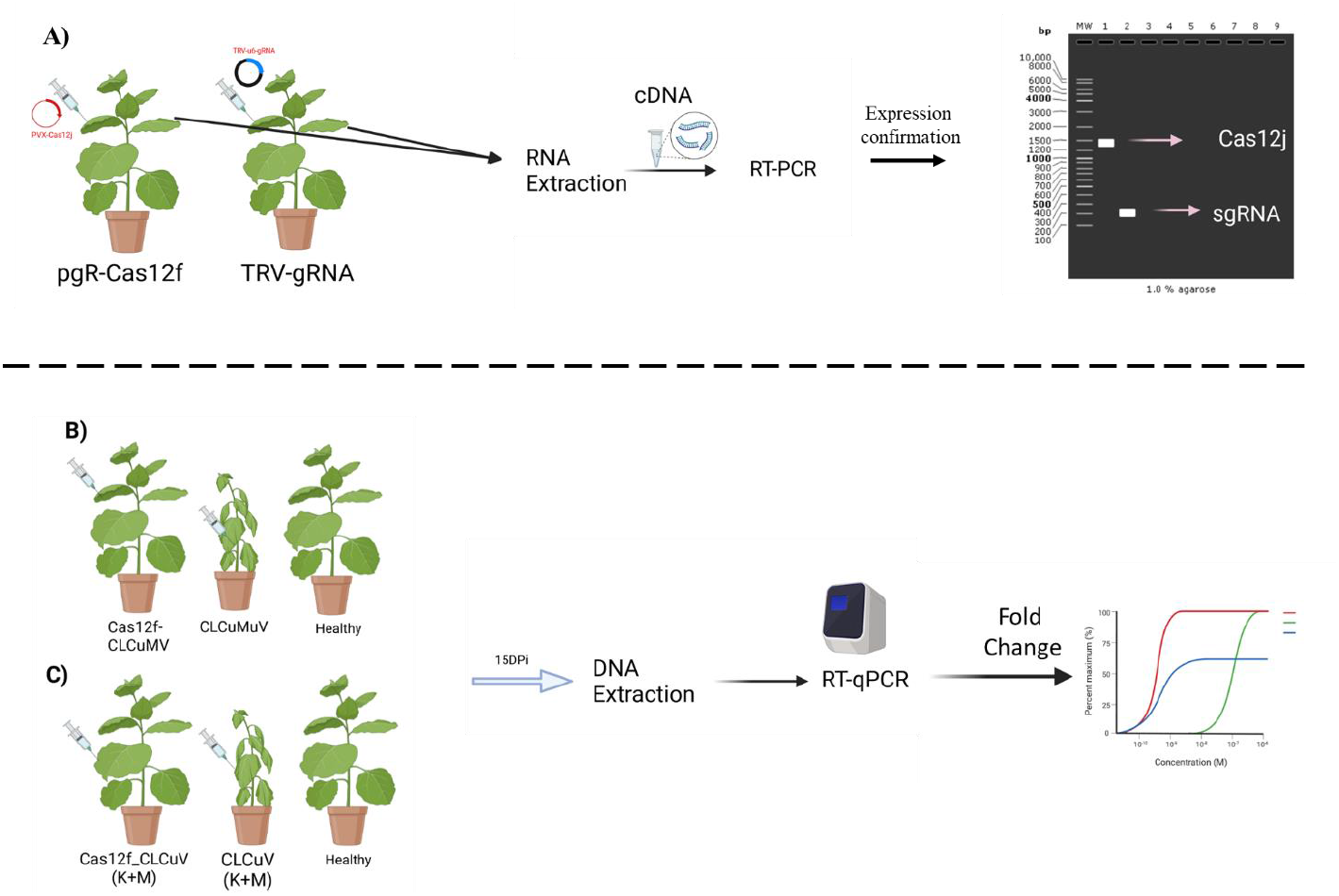
An overview of Agro-infiltration in *N. benthamiana*. **(A)** Transient expression analysis of Cas12f and sgRNA. pgR-Cas12f and TRV-gRNA constructs were inoculated in two weeks old *N. benthamiana* plants. 15d.p.i RNA was extracted and full-length expression of Cas12f and gRNA was confirmed with RT-PCR. **(B)** Left to right (1) Co-inoculation of Cas12f-gRNA complex with CLCuMuV (2) Co-inoculation of CLCuMuV only (3) Healthy control **(C)** Left to right (1) Co-inoculation of Cas12f-gRNA complex with CLCuV(K+M) (2) Co-inoculation of CLCuV(K+M) only (3) Healthy control. (B,C) DNA was extracted 15d.p.i and RT-qPCR was performed to identify the virus titer difference.

**Figure 3:**
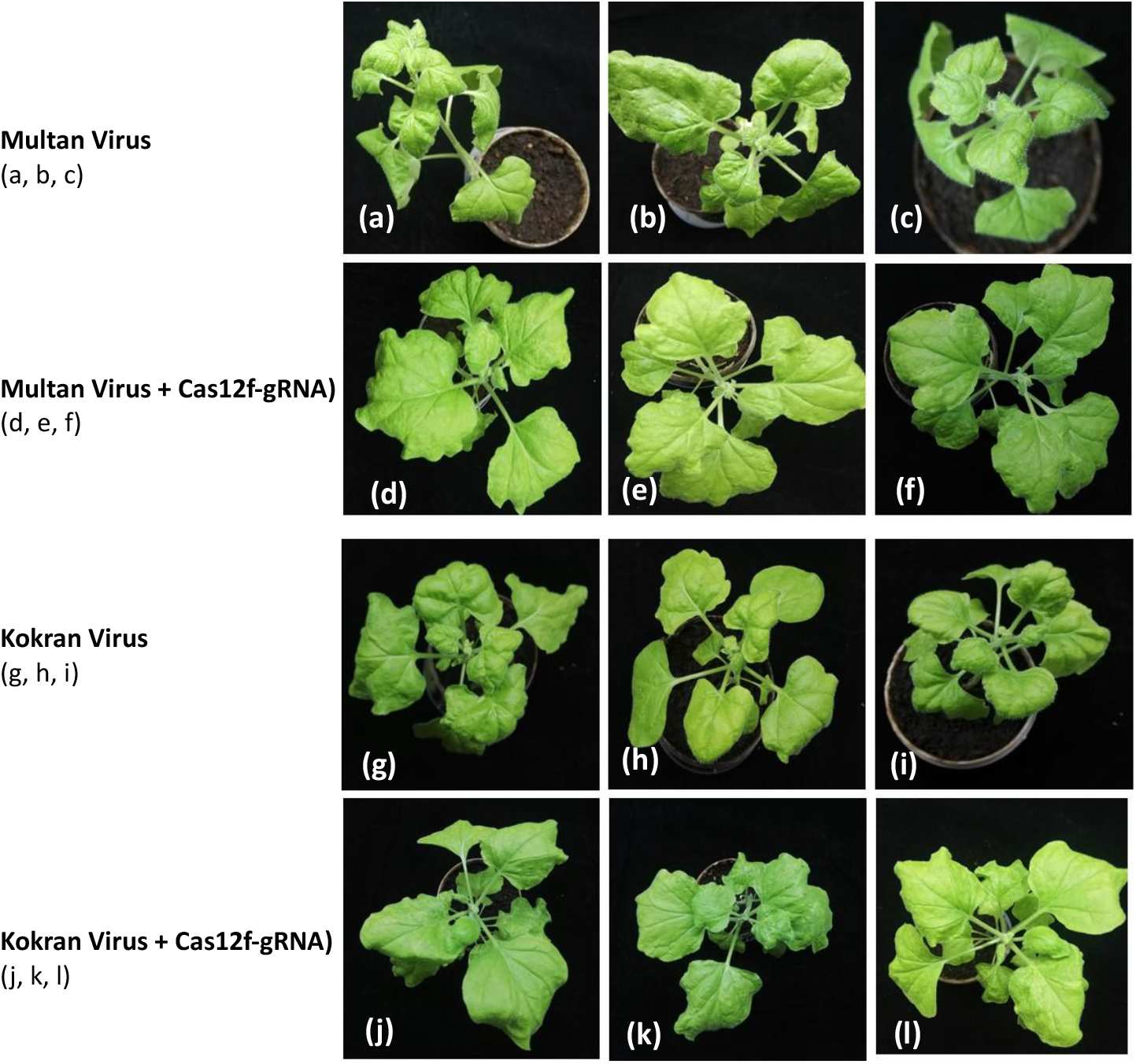
Transient expression analysis of Cas12f-gRNA against Multan virus and Kokran virus the N. benthamiana. **(a-c)** Control plants injected with only Multan virus showed higher leaf folding rate while **(d-f)** Plants were treated with Cas12f and gRNA followed by challenge with Multan virus showed minor symptoms **(g-i)** Control plants only injected with Kokran virus **(j-l)** Cas12f and gRNA expressed plants when challenged with Kokran virus showed no or less symptoms as compared to control ones

**Figure 3:**
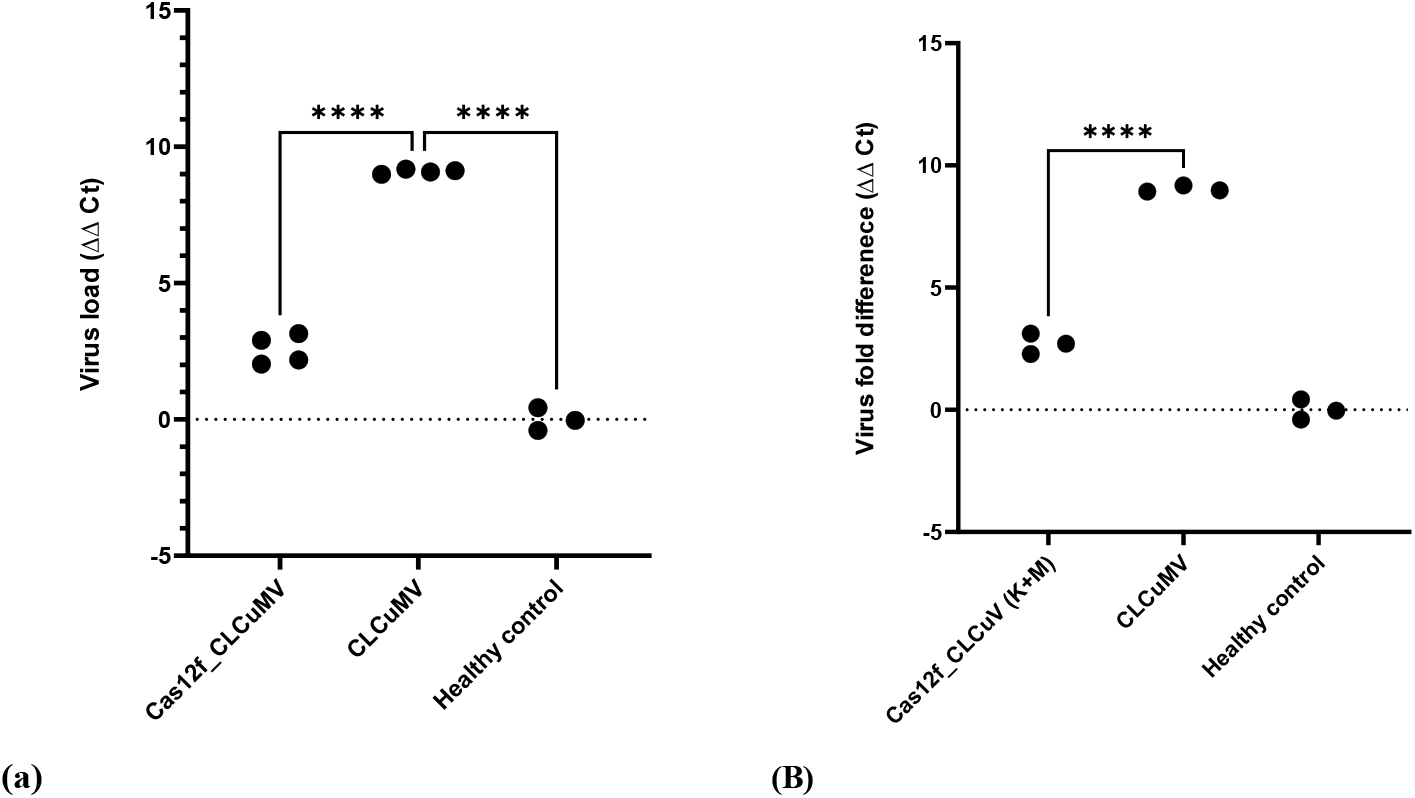
Determination of Virus titre by qPCR. The results were high resistance against CLCuMuV and CLCuKV. **(a)** It shows the virus (CLCuMuV) titre in the *N. benthamiana* with and without transient expression of Cas12f-gRNA complex. (Left to right) Cas12f_CLCuMV show the virus titer in plants with transient co-expression of Cas12f-gRNA complex, CLCuMuV shows the virus titre in plants without Cas12f-gRNA complex, Healthy control plants. Each dot present a seprate group of Plants used in the experiment. (**b)** It shows the combined virus (CLCuKV+ CLCuMuV) titer in the *N. benthamiana* with and without transient expression of Cas12f-gRNA complex. Cas12f_CLCuV (K+M) shows the virus titer in plants with transient co-expression of Cas12f-gRNA complex, CLCuV (K+M) shows the virus titer in plants without Cas12f-gRNA, Healthy control plants These results strongly support the idea that only one gRNA can be used to control broad-spectrum of ssDNA viruses.

**CLCuMuV**: Cotton leaf curl Multan virus
**CLCuV(K+M):** Mixture of Cotton leaf curl Kokran-virus and Cotton leaf curl Multan virus

Based on nonspecific cleavage activity of Cas12f we hypothesised that one gRNA should be enough to target more than one type of ssDNA viruses. For the proof of concept, we used a single gRNA (for CLCuMuV) against two major strains of begomovirus CLCuMuV and CLCuKV strain.

An inoculation mixture of CLCuMuV and CLCuKV was used along with pgR-Cas12f, TRV-gRNA. Surprisingly phenotype of these plants showed mild to no symptoms of CLCuD and single gRNA was effective against both CLCuMuV and CLCuKV.

To determine quantitative decrease in virus titer RT-qPCR was performed. DNA was isolated at 21 dpi from both plants treated with Cas12f (coupled with gRNA) and infected control plants (CLCuMuV, CLCuKV). Results showed highly significant decreaseof virus titter in plants harbouring Cas12f complex compared to control ones i.e., CLCuMuV alone and a mixture of CLCuMuV and CLCuKV.

Results shows first fully functional transient CRISPR-Cas12f system in plants. As a proof of concept, we also showed that Cas12f is much more robust and effective against ssDNA viruses as compared to Cas9 which only target dsDNA (Figure 3). Our data also show how plant RNA viral vectors can be used to study the effectivity of other Miniature Cas proteins and their targets.

## Material and Methods

### Identification of Begomovirus diversity

Begomoviruses show a very high mutation rate [21] and to identify the prevalence of current strain in the field multiple samples with symptoms of CLCuD (vein swelling, leaf curling) were collected from four major cotton-growing areas of the Punjab province of Pakistan during the year 2019. Genomic DNA was extracted and used as a template in PCR. Universal primers for begomoviruses, betasatellites and alphasatellites were used to amplify the genomic components [22]. Desired PCR products from infected samples were cloned in pTZ57R/T. After restriction digestion analysis, confirmed samples were sent for Sanger sequencing. The sequence reads were assembled and analysed for the detection of the sequences by using DNASTAR.

### gRNA designing

Based on high prevalence of CLCuMuV in Punjab region gRNA was designed on conserved region of pre-coat protein of CLCuMuV (Table 2). No PAM requirement for ssDNA make Cas12f-sgRNA more convenient in selecting the conserver region.

### Cloning and construct development

The pgR-Cas12f construct was developed for transient expression of Cas12f in the plant. For this purpose, Cas12f was amplified from vector pLBH545_Tet-Cas12f1 (Addgene plasmid # 112502) with primers containing restriction site Cla1 and Sal1 (Table 2). The confirmed PCR amplicon was ligated in pgR106 vector at Cla1 and Sal1 site to create pgR-Cas12f (Figure 1a). The pgR-Cas12f along with helper plasmid pSoup were transformed in *Agrobacterium tumefaciensstrain* strain GV3101 and confirmed colony was used for onward experiments.

For transient expression, tracrRNA-crRNA (gRNA-pc) against pre-coat protein of CLCuMuV was cloned under polymerase III promoter. For this purpose, a 263 bp long Cassette of gRNA-pc was synthesized with U6 terminator and Bbs1 restriction sites from Bio Basic Inc. (Table 2). The AtU6-26 promoter was amplified from pEN-Chimera and cloned in pTZ57R/T vector. Synthesized gRNA-pc was cloned under U6 promoter using BbsI site to develop a complete cassette composed of U6 promoter and gRNA-pc and U6 terminator. The U6-gRNA cassette was PCR amplified and cloned into pTRV-RNA2 at restriction site EcoR1 and Kpn1 and the resultant vector was named TRV2-gRNA (Figure 1). TRV2-gRNA with binary plasmid pTRV-RNA1 were transformed in *A. tumefaciens*, strain GV3101 (from now on combined refer as TRV-gRNA). The confirmed colony was used for the onward experiments. Infectious clones of Cotton leaf curl Multan virus (CLCuMuV) and Cotton leaf curl Kokran-virus (CLCuKV) used in this study are (Acc. No. KX656801) and (Acc. No. AM774295).

### Transient expression analysis of Cas12f and tracrRNA-crRNA

*Agrobacterium* cultures with plasmid pgR-Cas12f, TRV-gRNA were harvested, resuspended in infiltration media (10mM MgCl_2_) with final OD_600_ of 1.5. Different mixtures (Table 1) were prepared for inoculation. To activate the Vir genes acetosyringone was added and Cultures were kept in dark overnight. Nearly 2 weeks old *N. benthamina* plants were inoculated with these mixtures. Plants’ physiology was carefully noticed, and pictures were taken every three days.

The total RNA of inoculated plants was isolated after 15 d.p.i. (days post-infection) using TRIzol Reagent (Invitrogen) and then treated with DNaseI (Invitrogen) to remove any genomic DNA contamination. The cDNA was synthesized using RevertAid First Strand cDNA Synthesis Kit. RT-PCR was performed using Dream-Taq Green PCR master mix (Thermo Fisher). The RT-PCR was performed to validate the expression of full-length Cas12f and tracrRNA-crRNA in the plant. Furthermore, infectious clones CLCuMuV and CLCuKV were transformed in *A. tumefaciens* strain GV3101 to introduce infection in plants.

### Co-Inoculation and analysis of virus titter

#### Specific immunity

Initially, we evaluated the effectiveness of Cas12f (coupled with gRNA) against plant CLCuMuV. For this purpose, *Agrobacterium* cultures containing pgR-Cas12f, TRV-gRNA, and CLCuMuV were inoculated in *N. benthamina* plants (Table 1). Plants’ physiology was carefully recorded. Plant DNA was isolated at 15 d.p.i. using CTAB method [19]. A SYBR^®^Green PCR Master Mix (Thermo Scientific) was used for qPCR-based analysis to quantify the virus titter.

#### Wide-ranging immunity

To evaluate broad-spectrum protection of activated Cas12f, an inoculation mixture of CLCuMuV and CLCuKV was used (table2). Plants’ physiology was carefully recorded. DNA was isolated at 15 d.p.i. using CTAB method [19]. A SYBR^®^Green PCR Master Mix (Thermo Scientific) was used for qPCR-based analysis to quantify the virus titter.

## Discussion

Plant DNA viruses are of great economic importance and responsible for great loss every year. Begomovirus is a group of DNA viruses, responsible for Cotton leaf curl Disease (CLCuD). It is a major constrain on cotton yield in Pakistan and India. Because of high diversity and mutation rate in disease complex it is very difficult to design a single strategy against the disease complex [1–3, 8, 13, 16, 21, 22]. CRISPR/Cas system has been proved as one of the most powerful tools in molecular biology for in vivo target-specific editing and manipulation to genomic DNA of both animals and plants. CRISPR/Cas9 system has been widely utilized to develop resistance in plants against viruses. This system can either be utilized to target plant viruses directly or can be employed to alter susceptible genes in plants to confer resistance [9,23–26]

The viruses against which CRISPR/Cas9 system has been utilized to induce resistance in plants includes beet severe curly top virus (BSCTV), cotton leaf curl virus (CLCuV), bean yellow dwarf virus (BeYDV) and tomato yellow leaf curl virus (TYLCV) in *N. bentiamana* and *Arabidopsis thaliana* plants. Cleavage site generated by Cas9 is repaired by NHEJ, this may lead to adaptation of viruses according to spacer or protospacer associated motif (PAM) which is an essential part for Cas9 to work [27–29]. Another characteristic limitation of Cas9 is that it is unable to target RNA viruses directly. To overcome this shortcoming, it is suggested to employ CRISPR-Cas13a along with CRISPR-Cas9. LwaCas13a system is reported to have ability to express five distinct gRNAs and hence can target five sites simultaneously [30].However, the previous Cas proteins i.e. Cas9 and Cas 13a are efficient in targeted cleavage of dsDNA and ssRNA but their activity is highly dependent on protospacer adjacent motif (PAM). Secondly, Cas9 (100-200 kDa) and Cas13a (~168 kDa) are large proteins which limits their use widely [31]. In contrast another Cas protein, Cas12f has size of 40-70 kDa and one of the variants, Cas 14a, which we used in our study has ability to cleave ssDNA with high efficiency and independent of PAM-sequence so there is nearly no chance for virus to adopt to PAM sequence and by pass Cas12f activity [14].

We were able to establish first fully functional transient miniature CRISPR-Cas system. RNA viruses have proved to be instrumental tools in robust transient expression system. For the expression of foreign gene PVX shows a little more capacity of~1.6kb as compared to TRV vector ~ 600bp. Based on their size Cas protein was cloned in PVX while gRNA cassette was cloned in TRV binary vector. While both PVX and TRV showed negligible symptoms on plant but were highly efficient to produce a systematic expression of corresponding transcript.

In the current study, the efficiency of CRISPR/Cas12f system has been investigated to induce broad-spectrum resistance in *N. benthamiana* against two most prevalent strains of CLCuV in Pakistan i.e. CLCuMuV and CLCuKV. The sgRNA was designed against pre-coat (pc) protein of CLCuMuV. Plasmids containing CRISPR/Cas12f, and gRNA were transiently expressed in *N. benthamiana* and were challenged with infectious clone of CLCuMuV and CLCuKV.

Cas12f showed effecient cleavage of both the viruses using only one gRNA. plants showed only minor curling at edges of the leaves as compare to infected plants and a visible difference was obvious between the health and growth of Treated and infected plants. Surprisingly, qPCR results revealed multifold (8-10 time) less virus titer on average than infected plants. This high resistance and surprising results may have been a result of independency of Cas12f on PAM sequences against ssDNA particles. These promising results have revealed that CRISPR/Cas12f system can be utilized to induce broad-spectrum viral resistance in plants without fear of playing any role in viral diversity in short time.

## Supporting information

arranged qPCR samples and analysed

Graph Pad file for Multan and kokran virus titer

